# In depth analysis of Cyprus-specific mutations of SARS-CoV-2 strains using computational approaches

**DOI:** 10.1101/2021.06.08.447477

**Authors:** Anastasis Oulas, Jan Richter, Maria Zanti, Marios Tomazou, Kyriaki Michailidou, Kyproula Christodoulou, Christina Christodoulou, George M. Spyrou

**Affiliations:** Bioinformatics Department, The Cyprus Institute of Neurology & Genetics, Nicosia, Cyprus; Molecular Virology Department, Cyprus Institute of Neurology and Genetics, Nicosia, Cyprus; Biostatistics Unit, The Cyprus Institute of Neurology and Genetics, Nicosia, Cyprus; Neurogenetics Department, Cyprus Institute of Neurology & Genetics, 6 Iroon Avenue, 2371; The Cyprus School of Molecular Medicine, Nicosia, Cyprus

## Abstract

This study aims to characterize SARS-CoV-2 mutations which are primarily prevalent in the Cypriot population. Moreover, using computational approaches, we assess whether these mutations are associated with changes in viral virulence. We utilize genetic data from 144 sequences of SARS-CoV-2 strains from the Cypriot population obtained between March 2020 and January 2021, as well as all data available from GISAID. We combine this with countries’ regional information, such as deaths and cases per million, as well as COVID-19-related public health austerity measure response times. Initial indications of selective advantage of Cyprus-specific mutations are obtained by mutation tracking analysis. This entails calculating specific mutation frequencies within the Cypriot population and comparing these with their prevalence world-wide throughout the course of the pandemic. We further make use of linear regression models to extrapolate additional information that may be missed through standard statistical analysis. We report a single mutation found in the *ORF1ab* gene (S6059F) that appears to be significantly enriched within the Cypriot population. We further analyse this mutation using regression models to investigate possible associations with increased deaths and cases per million. Moreover, protein structure prediction tools show that the mutation infers a conformational change to the protein that significantly alters its structure when compared to the reference protein. Investigating Cyprus-specific mutations for SARS-CoV-2 can not only lead to important findings from which to battle the pandemic on a national level, but also provide insights into viral virulence worldwide.

## Introduction

Cyprus’ first case of COVID-19 was reported on March 9^th^ 2020, with a gradual increase after that to formulate the first peak of the virus transmission seen in late March with all major Cypriot cities affected by the virus. This first wave was relatively mild with respect to reported daily new cases and deaths reaching a maximum of 58 cases on April 1^st^ and 2 deaths. The second wave of virus transmissions hit Cyprus in mid-October and gradually increased to peak in December with a maximum of 907 and 8 reported daily new cases and deaths, respectively. This summary of the SARS-CoV-2 spread in the Cypriot population dictates that Cyprus is one of the least affected European countries during this pandemic (mostly with respect to deaths per million). This can be attributed to the rapid response time for austerity measures and the effective quarantine process for confirmed cases and their contacts. In addition, Cyprus is ranked 3^rd^ in Europe as for the number of COVID tests performed per million (>2,4M tests).

Multiple national studies have been undertaken by a plethora of countries throughout the world, in order to generate and analyse high throughput sequencing country-specific data for SARs-CoV-2 strains [1–8]. Recently, the Cyprus Institute of Neurology and Genetics (CING) also published a study based on 144 NGS samples obtained from the Cypriot population representing the first documented genomic and epidemiological characterization of these samples [9]. In this study we expand on this work, by initially performing basic lineage analysis, as previously reported [9], to identify the dominant SARS-CoV-2 lineage(s) in Cyprus as well as a phylogenetic tree analysis to map the Cypriot strains against other strains present throughout the world. We further perform a more thorough, in depth genomic analysis of these samples by implementing variant-calling and displaying an overview of the genomic variation and reported frequencies in the Cypriot strains. We then focus on the 9 spike (S) protein mutations that delineate the UK B.1.1.7 lineage and how their founder effect in Cyprus has impacted the number of cases and deaths in the population. Furthermore, we identify Cypriot-specific/dominant mutations and investigate them using mutation tracking analysis in order to isolate their origin, determine their founder effect in Cyprus and also trace their overall prevalence and propagation in other countries. We capitalize on our previously published generalized linear models [10] to undertake virulence analysis on Cypriot-dominant mutations and show how their presence has affected the number of cases and deaths within the country. Finally, we perform structural modelling of the alternate versus the reference mutation for selected mutations of interest and view their effects at the viral protein level.

## Materials and Methods

### Raw data analysis

The Burrows-Wheeler Aligner (BWA) [11], version: 0.7.15 was used to map the raw reads to Wuhan-Hu-1 (NCBI ID:NC_045512.2). Duplicate reads, which are likely to be the results of PCR bias, were marked using Picard (http://broadinstitute.github.io/picard/) version: 2.6.0. SAMtools [12], version: 0.1.19, was used for additional BAM/SAM file manipulations. The Genome Analysis Tool Kit (GATK) [13], version 3.6.0, HaplotypeCaller method was used for single nucleotide polymorphism (SNP) and insertion/deletion (indel) variant calling generating vcf files. All mutations were annotated according to the reference Wuhan-Hu-1 strain. These annotations include: genomic position, reference (REF) genotype, alternate (ALT) genotype, gene, encoded amino-acid with REF genotype, encoded amino-acid change (if not synonymous) with ALT genotype, amino-acid position according to main encoded viral protein (annotation file available as Supplementary Table S1). Finally, the GATK FastaAlternateReferenceMaker method was used for consensus sequence extraction from the vcf files.

### Lineage assignment

The consensus sequences of all 144 Cypriot strains were uploaded to the Pangolin COVID-19 lineage assigner interface [14] (https://pangolin.cog-uk.io/). Further analysis of results and generation of visualizations was performed in R V.3.6.1 [15].

### Phylogenetic analysis and comparison with other strains

Full genomic SARS-CoV-2 sequences of high sequencing resolution were obtained from GISAID (https://www.gisaid.org/, last accessed 15/01/21). The nextstrain [16] pipeline was downloaded locally and the commands for filtering, aligning and constructing phylogeny were used according to the nextstrain’s best practices. MAFFT [17] was used to construct a multiple sequence alignment (MSA). Phylogeny was estimated using the RAxML [18] maximum likelihood algorithm for phylogenetic tree construction. Variant calling for the GISAID strains was achieved using the snp_sites tool available through github (https://github.com/sanger-pathogens/snp-sites).

### Mutation tracking analysis

Relative frequencies across countries with at least one occurrence of the selected mutations of interest was visualized as bar plots across time (months). This provides an indication of the spread of the studied mutations across the general population. Analysis was performed using R (packages: dplyr, tidyr, ggplot2, ggtree, phytools, phangorn).

### Identifying Cyprus-specific mutations

A generalized linear model was applied (see formula 1 below), whereby the absence or, presence (values 0 or 1 respectively) of the S6059F mutation (*mut* variable) was assessed across samples from Cyprus, as well as the rest of the world (as defined by the *classes* variable, 1 denoting Cyprus and 0 for other countries).

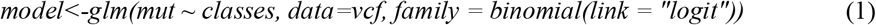

An *odds* ratio analysis was also performed using a simple formula (see 2 below). Whereby, *cycounts* denotes the number of times the S6059F mutation was observed in Cyprus (0 for absence, 1 for presence) and similarly *othercounts* denotes the number of times the mutation was observed in other countries.

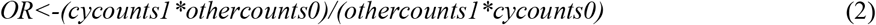

### Structural Analysis

Comparative structural modelling was carried out for unknown protein structures using the template-based web server, I-TASSER [19]. The accuracy of the method was assessed based on the I-TASSER template modeling score (TM-score), which indicates the structural similarity between model and templates. A TM-score higher than 0.5 indicates a model of correct topology, while a TM-score of less than 0.17 means a random topology [19]. I-TASSER was selected for protein structure modelling, since it outperformed other servers according to results from the 14th Community Wide Experiment on the Critical Assessment of Techniques for Protein Structure Prediction (CASP14) (https://zhanglab.ccmb.med.umich.edu/casp14/, last accessed 23/03/2021). Mutagenesis was performed using the DynaMut suite (http://biosig.unimelb.edu.au/dynamut/prediction, last accessed 23/03/2021). The PyMOL software (v0.99) was used for the visualization of the protein molecules.

Relative solvent accessibility (RSA) was calculated using the Missense3D algorithm [20]. The DynaMut webserver [21], was used to visualize non-covalent molecular interactions, calculated by the Arpeggio algorithm [22]. Protein-protein complexes were constructed using the ClusPro (v2.0) [23] and HDOCK [24] algorithms and binding affinities and dissociation constants (Kd) were calculated using the PRODIGY webserver [25]. RNA-protein docking simulations were carried out using the HDOCK [31] and MPRDock [32] algorithms. For RNA-protein docking simulations, as active residues we selected the active site residues of the SARS-CoV NSP14 protein, since the two proteins share a 99.1% sequence similarity [26]. Structural alignment was performed using the align tool of PyMOL and all-atom RMSD values were calculated without any outliers’ rejection, with zero cycles of refinement. All docking simulations were performed in triplicates.

## Results

### Dominant lineages present throughout the COVID-19 pandemic in Cyprus – emphasis on UK lineage B.1.1.7

One of the most widely used international systems for detecting lineages that contribute most to active spread is the dynamic nomenclature system presented by Rambaut et al [14]. Lineage analysis of the 144 Cypriot viral strain genomes showed 16 major lineages including strains originating from both A and B lineages, which are denoted as the root lineages of the phylogeny of SARS-CoV-2 (**Figure 1**). Dominant lineages included: (i) B.1.258 (51.03% of sequenced strains), with most common countries of origin being UK, Denmark and Czech Republic and (ii) the UK lineage, B.1.1.7, having drawn attention by the recent outbreak in the UK with reported increased rates of viral transmission [27–29], which was detected with high prevalence within the Cypriot population (13.1% of analysed strains). This lineage is characterized by a series of 9 mutations (deletion HV69-70, deletion Y144, N501Y, A570D, P681H, T716I, S982A, D1118H, D614G) within the viral S protein. The D614G mutation has now become dominant in all strains but was included here for completion. Mutation tracking for 7/9 of these mutations reveals that the specific lineage was firstly reported in Cyprus on December 2020 and was a consequence of Cypriot citizens returning to Cyprus after visiting the UK. Moreover, the founder effect of this lineage within the Cypriot population appears to be associated with an increase in number of cases, a phenomenon also reported for the UK [29] (see **Figure 2**).

**Figure 1.**
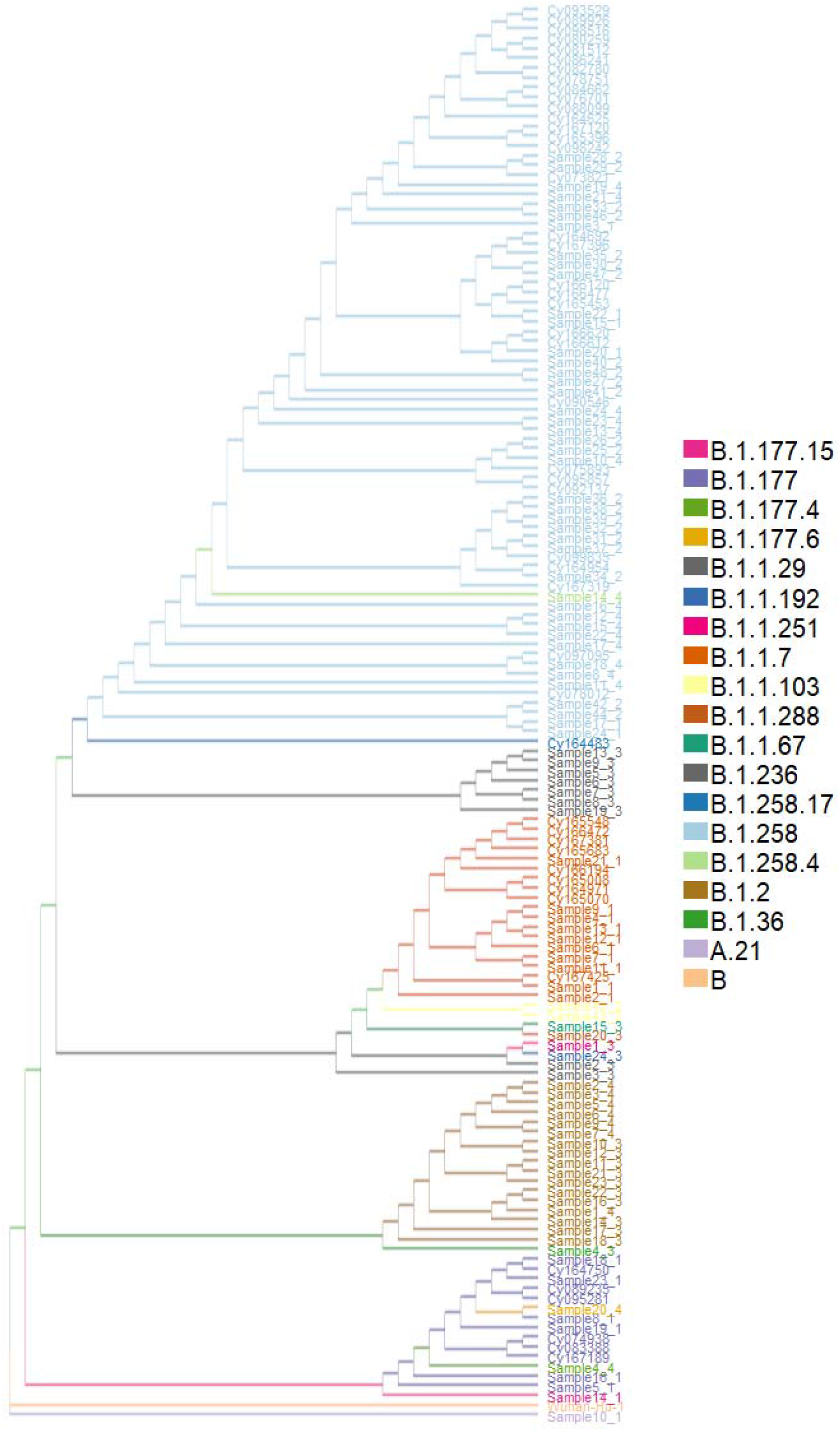
Phylogenetic tree of Cyprus-derived strains from COVID samples and lineage analysis using Pangolin lineage analysis tool.

**Figure 2.**
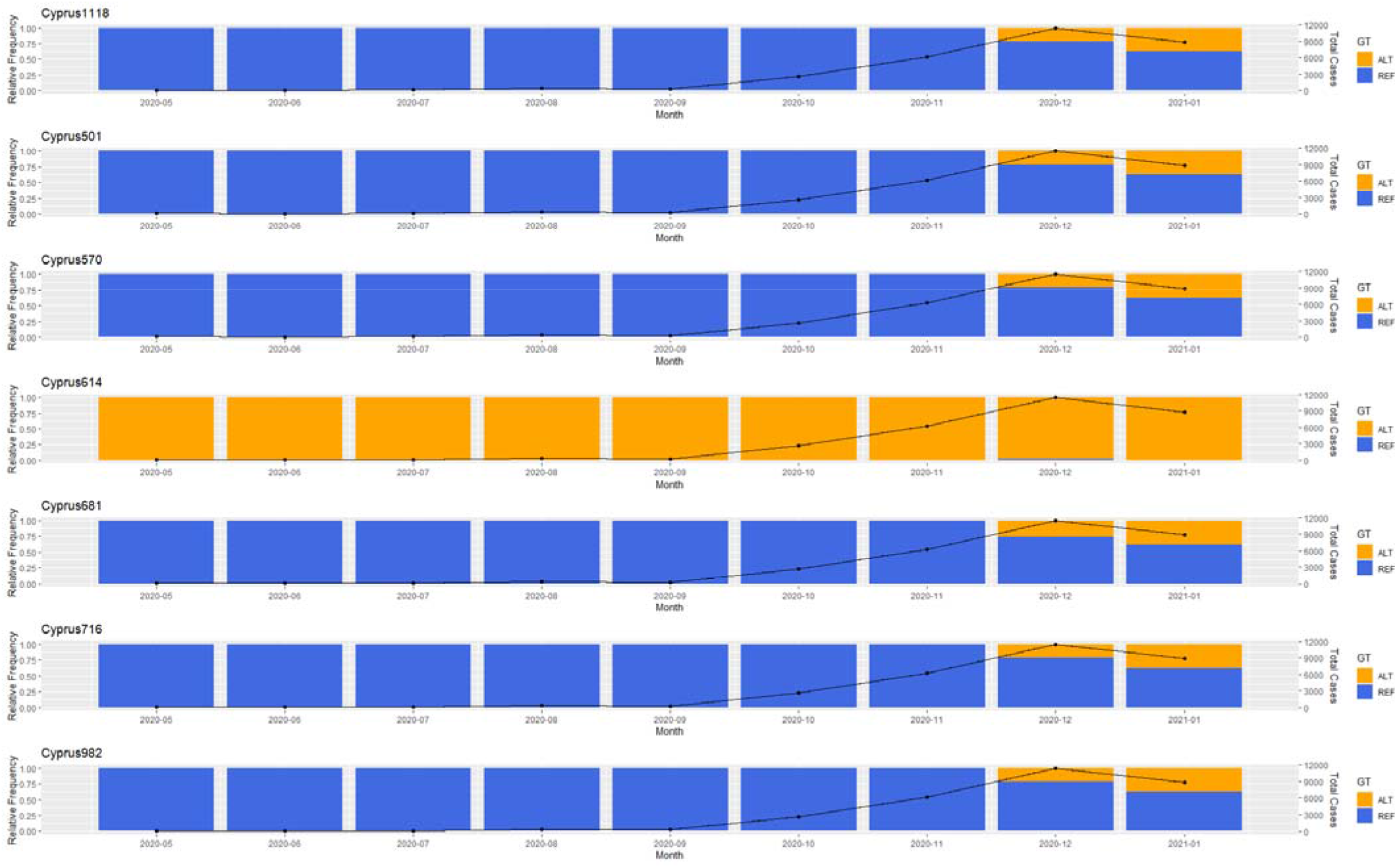
Mutation tracking of 7/9 mutations that delineate the B.1.1.7 lineage originating from the UK. The alternate (ALT) form of the mutations is shown in orange while the reference (REF) form of the mutations are denoted in blue. The black line shows the number of cases for each country per month.

### Cyprus specific mutations

Cyprus-specific mutations were obtained by counting the frequency of all single nucleotide mutations present within the vcf file generated from our 144 sequenced samples. We selected for mutations that appeared in at least 30% of the samples and thus identified 18 mutations that met this criterion. These included mutations that were very cosmopolitan (e.g. D614G mutation) but also included mutations that appeared primarily in the Cypriot population according to our sequenced samples. We performed an odds ratio (OR) analysis to investigate which mutations appear to be more frequent in the Cypriot population vs. strains obtained from the rest of the word. We also applied a generalized linear model (glm) with a logistic regression using the *logit* function in order to obtain a model for each of the 18 selected mutations. The model assumes a value of 1 if the mutation is present and 0 if not, through this way trying to see how well this fits the ideal situation of mutations only occurring in the Cypriot samples vs. the rest of the world (for details see materials and methods). P-values were generated to show how well the glm performs under this scenario. We specifically highlight the S6059F mutation, located in the NSP14 protein of the *ORF1ab* gene, because it obtained the highest OR statistic (6921.17) and was also deemed most significant according to our glm model (p-value = 6.94E-180) (see **Table 1**). We performed mutation tracking analysis to further investigate the prevalence of this specific mutation throughout the course of the pandemic, both in Cyprus as well as worldwide. The founder effect appeared to have occurred in Russia, as shown by timestamps for the strains as well as phylogenetic tree analysis for the specific strains with the S6059F mutation (see **Figure 3**). This event did not lead to the establishment of this mutation as the dominant form within the Russian population. Similar results appear for other countries with at least one reported strain with this mutation. This is in contrast to Cyprus where the alternate S6059F mutation clearly appears to be replacing the reference form (**see Figure 4**).

**Table 1.**
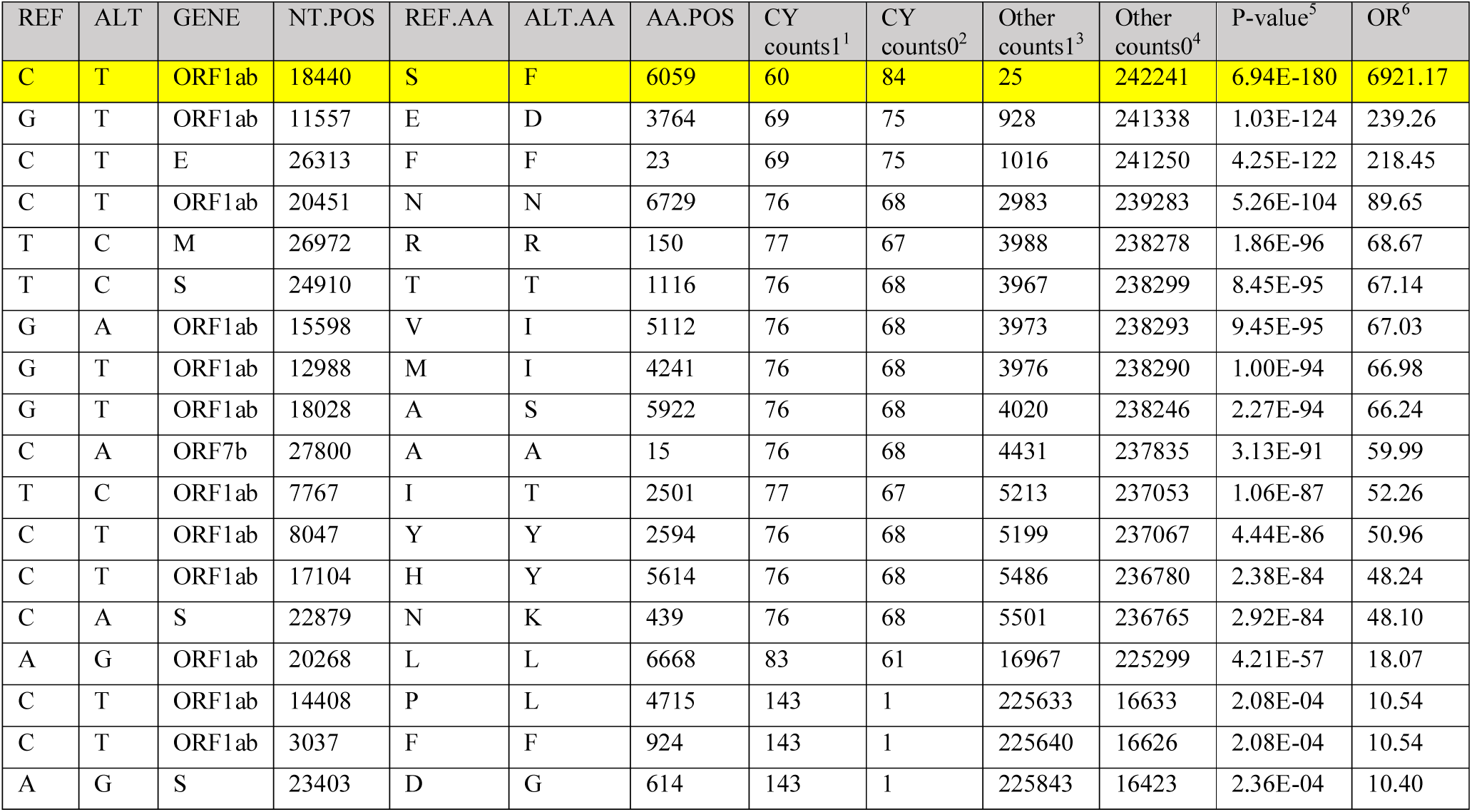
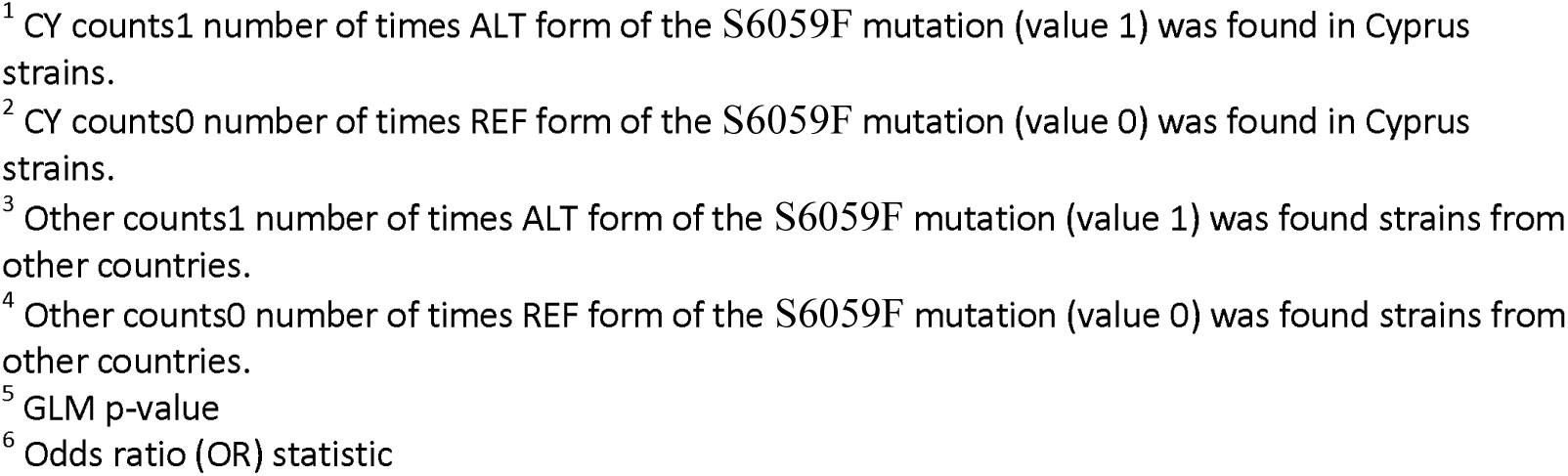
Cyprus specific mutations ordered by p-value for best fit according to our glm model.

| REF | ALT | GENE | NT.POS | REF.AA | ALT.AA | AA.POS | CY counts1<sup>1</sup> | CY counts0<sup>2</sup> | Other counts1<sup>3</sup> | Other counts0<sup>4</sup> | P-value<sup>5</sup> | OR<sup>6</sup> |
| --- | --- | --- | --- | --- | --- | --- | --- | --- | --- | --- | --- | --- |
| C | T | ORF1ab | 18440 | S | F | 6059 | 60 | 84 | 25 | 242241 | 6.94E-180 | 6921.17 |
| G | T | ORF1ab | 11557 | E | D | 3764 | 69 | 75 | 928 | 241338 | 1.03E-124 | 239.26 |
| C | T | E | 26313 | F | F | 23 | 69 | 75 | 1016 | 241250 | 4.25E-122 | 218.45 |
| C | T | ORF1ab | 20451 | N | N | 6729 | 76 | 68 | 2983 | 239283 | 5.26E-104 | 89.65 |
| T | C | M | 26972 | R | R | 150 | 77 | 67 | 3988 | 238278 | 1.86E-96 | 68.67 |
| T | C | S | 24910 | T | T | 1116 | 76 | 68 | 3967 | 238299 | 8.45E-95 | 67.14 |
| G | A | ORF1ab | 15598 | V | I | 5112 | 76 | 68 | 3973 | 238293 | 9.45E-95 | 67.03 |
| G | T | ORF1ab | 12988 | M | I | 4241 | 76 | 68 | 3976 | 238290 | 1.00E-94 | 66.98 |
| G | T | ORF1ab | 18028 | A | S | 5922 | 76 | 68 | 4020 | 238246 | 2.27E-94 | 66.24 |
| C | A | ORF7b | 27800 | A | A | 15 | 76 | 68 | 4431 | 237835 | 3.13E-91 | 59.99 |
| T | C | ORF1ab | 7767 | I | T | 2501 | 77 | 67 | 5213 | 237053 | 1.06E-87 | 52.26 |
| C | T | ORF1ab | 8047 | Y | Y | 2594 | 76 | 68 | 5199 | 237067 | 4.44E-86 | 50.96 |
| C | T | ORF1ab | 17104 | H | Y | 5614 | 76 | 68 | 5486 | 236780 | 2.38E-84 | 48.24 |
| C | A | S | 22879 | N | K | 439 | 76 | 68 | 5501 | 236765 | 2.92E-84 | 48.10 |
| A | G | ORF1ab | 20268 | L | L | 6668 | 83 | 61 | 16967 | 225299 | 4.21E-57 | 18.07 |
| C | T | ORF1ab | 14408 | P | L | 4715 | 143 | 1 | 225633 | 16633 | 2.08E-04 | 10.54 |
| C | T | ORF1ab | 3037 | F | F | 924 | 143 | 1 | 225640 | 16626 | 2.08E-04 | 10.54 |
| A | G | S | 23403 | D | G | 614 | 143 | 1 | 225843 | 16423 | 2.36E-04 | 10.40 |

<sup>1</sup> CY counts1 number of times ALT form of the S6059F mutation (value 1) was found in Cyprus strains.
<sup>2</sup> CY counts0 number of times REF form of the S6059F mutation (value 0) was found in Cyprus strains.
<sup>3</sup> Other counts1 number of times ALT form of the S6059F mutation (value 1) was found strains from other countries.
<sup>4</sup> Other counts0 number of times REF form of the S6059F mutation (value 0) was found strains from other countries.
<sup>5</sup> GLM p-value
<sup>6</sup> Odds ratio (OR) statistic

**Figure 3.**
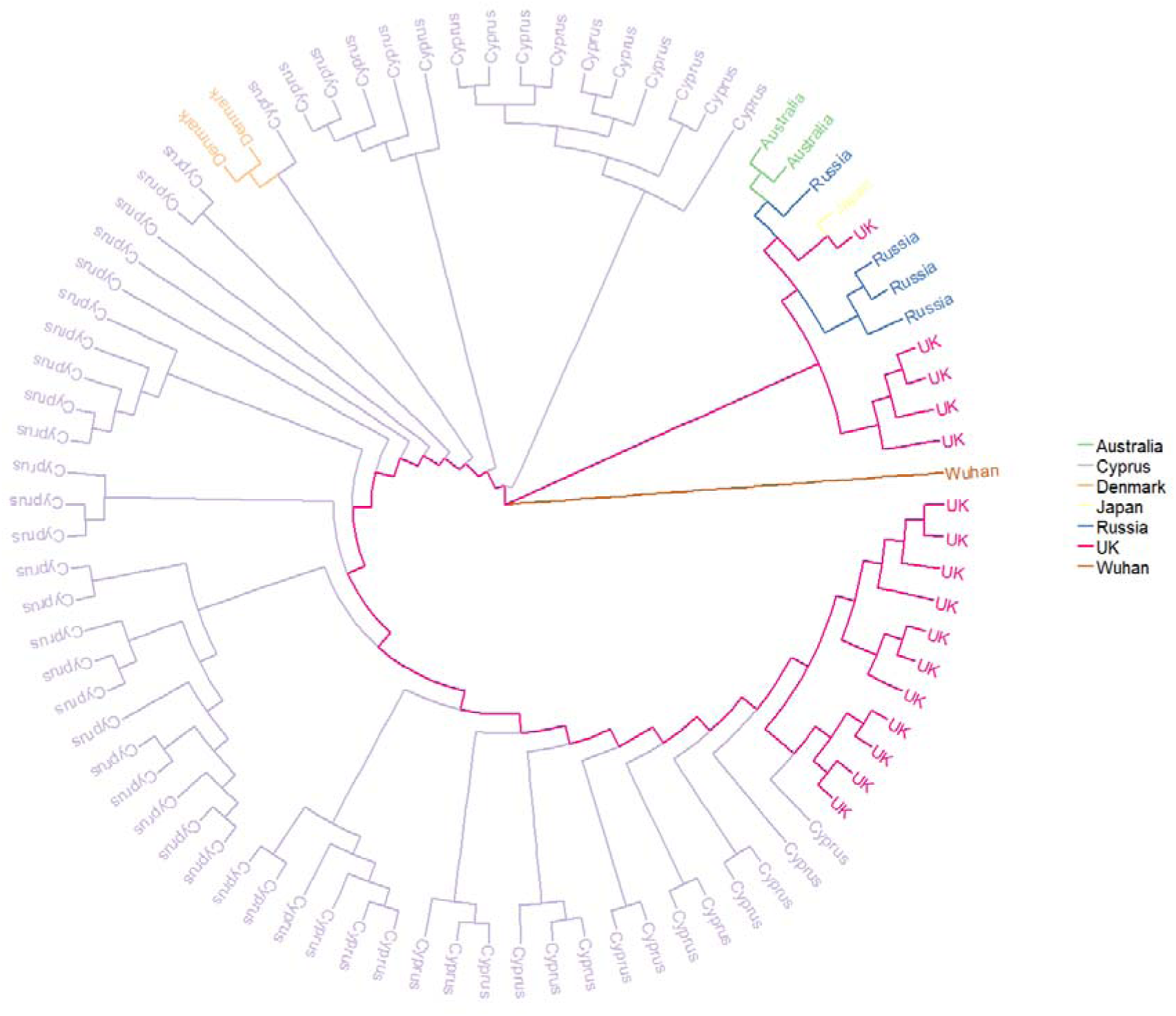
Phylogenetic tree of strains with the S6059F mutation labelled by country. The reference Wuhan strain has been added for completion.

**Figure 4.**
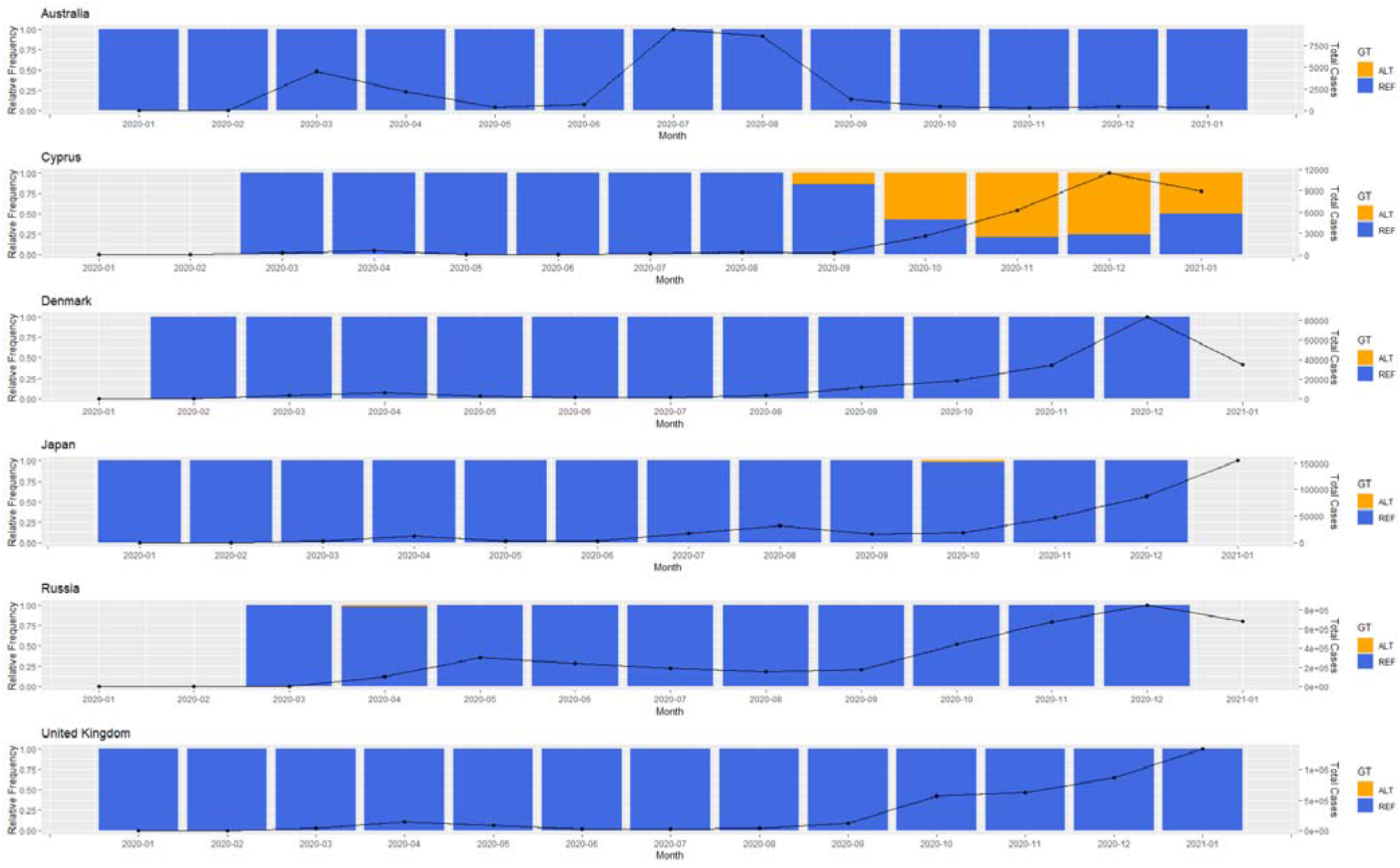
Mutation tracking of the S6059F mutation in Cyprus and other countries with at least one reported strain with the specific mutation. The alternate (ALT) and reference (REF) forms of the mutations are denoted in orange and blue respectively. The black line shows the number of cases for each country on a monthly basis.

The NSP14 protein is bifunctional and contains two domains: a 3′-to-5′ exoribonuclease (ExoN) and a guanine-N7-methyltransferase (N7-MTase). It is presumed that the N7-MTase domain supports mRNA capping, while the ExoN domain is believed to mediate proofreading during genome replication [30]. Previous studies have shown that ExoN knockout mutants or severe acute respiratory syndrome coronavirus (SARS-CoV) exhibit a dysfunctional yet viable hypermutation phenotype both in cell culture as well as animal models [31–33] while a SARS-CoV-2 ExoN knockout mutant was found to be unable to replicate [30].

In order to investigate for proofreading dysfunctionality in the strains with the S6059F mutation within the Cypriot population, we compared mutation counts for all strains with the alternative (ALT) form of this mutation to the strains with the reference (REF) form. The 60 strains with the ALT S6059F mutation all belonged to the B.1.258 lineage and showed a significantly higher mean for the number of mutations when compared to the REF S6059F strains (Wilcoxon test - p-value 0.00062 – see **Figure 5**). We propose that this difference in mutation counts within the strains with different forms of the S6059F mutation (ALT vs. REF) can be a attributed to increased hypermutability as a consequence of the amino acid change in the NSP14 protein.

**Figure 5.**
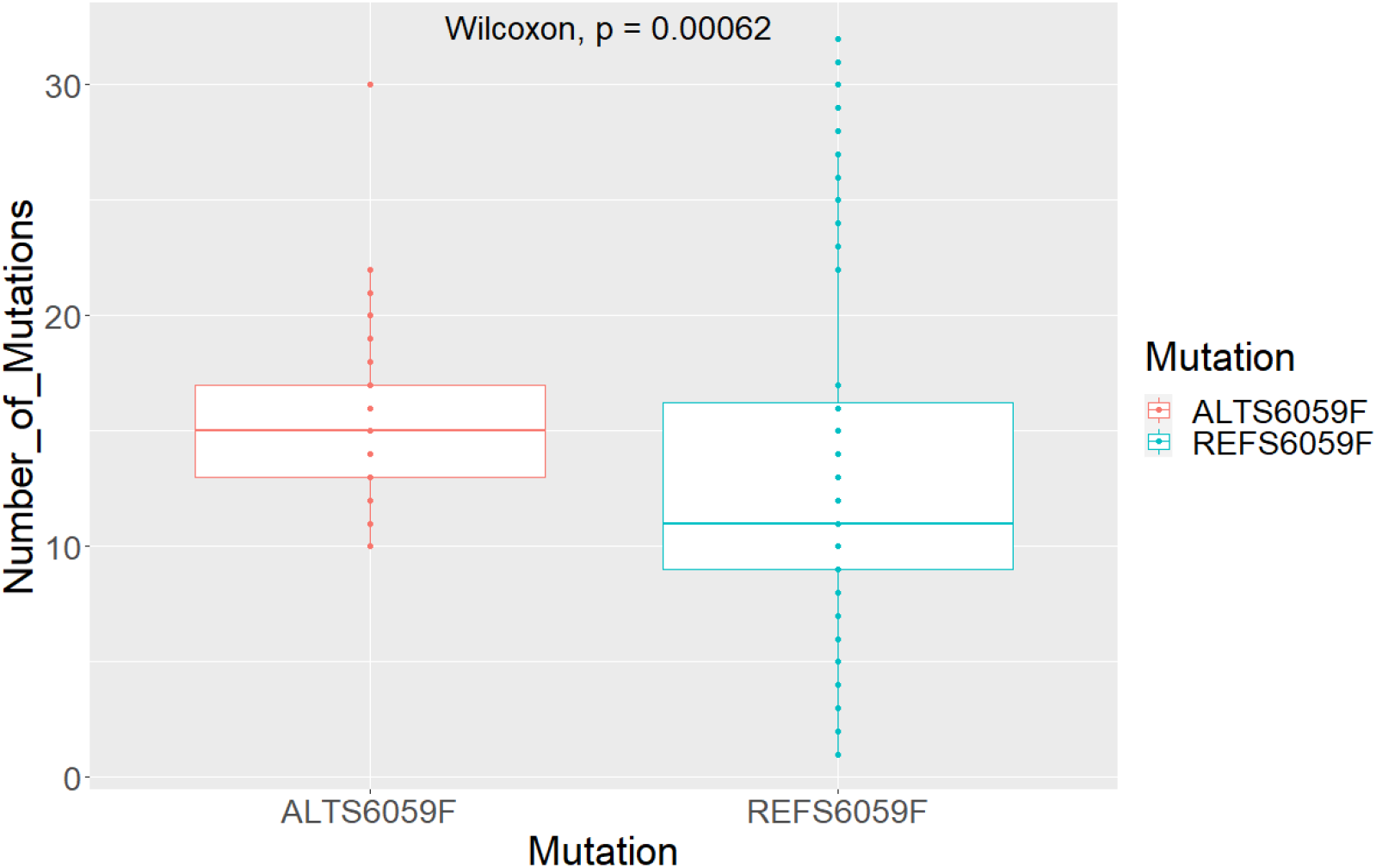
Number of mutations reported in strains with and without the S6059F mutation within the Cypriot population.

### Virulence analysis of Cyprus-specific SARS-COV-2 mutations

Capitalizing on previously published generalized linear models (glms) that provide a measure of association between specific SARs-CoV-2 mutations and increased or decreased viral cases or deaths [10], we proceeded to perform virulence analysis on the 18 mutations found to be prevalent in the Cypriot population. We focused on the S6059F mutation due to its exclusive propagation within Cyprus compared to the rest of the world. Our glms are uniquely designed to incorporate mutation frequency, austerity measure response time and mutation occurrence information as predictor variables and cases or deaths per million as response variables into a single model. This allows for a fit that determines whether a given mutation is positively or negatively correlated with number of cases and deaths per country.

We focused on geographic regional data reporting deaths and transmissions according to the Worldometers.info website (last accessed 08/03/21). Populations harbouring the mutation show a higher mean of deaths per million (1363) compared to the converse (1206). Statistical analysis of the populations with and without the S6059F mutation shows substantial evidence that the two distributions are significantly different (Wilcoxon test – *p*-value 2.2e-16) (see **Figure 6A**). Results obtained from using the cases per million parameter to assess the different distributions, show a reversed pattern with samples with the mutation exhibiting a lower mean of cases (49224) compared to the reference samples (64630) (see **Figure 6B**). Statistical analysis further provides evidence that the two distributions are significantly different (Wilcoxon test – *p*-value 2.2e-16). We also included response time separation in the box plots which shows how the mutation segregates across countries that responded differently to the COVID austerity measures (see supplementary **Figure S1A,B**).

**Figure 6.**
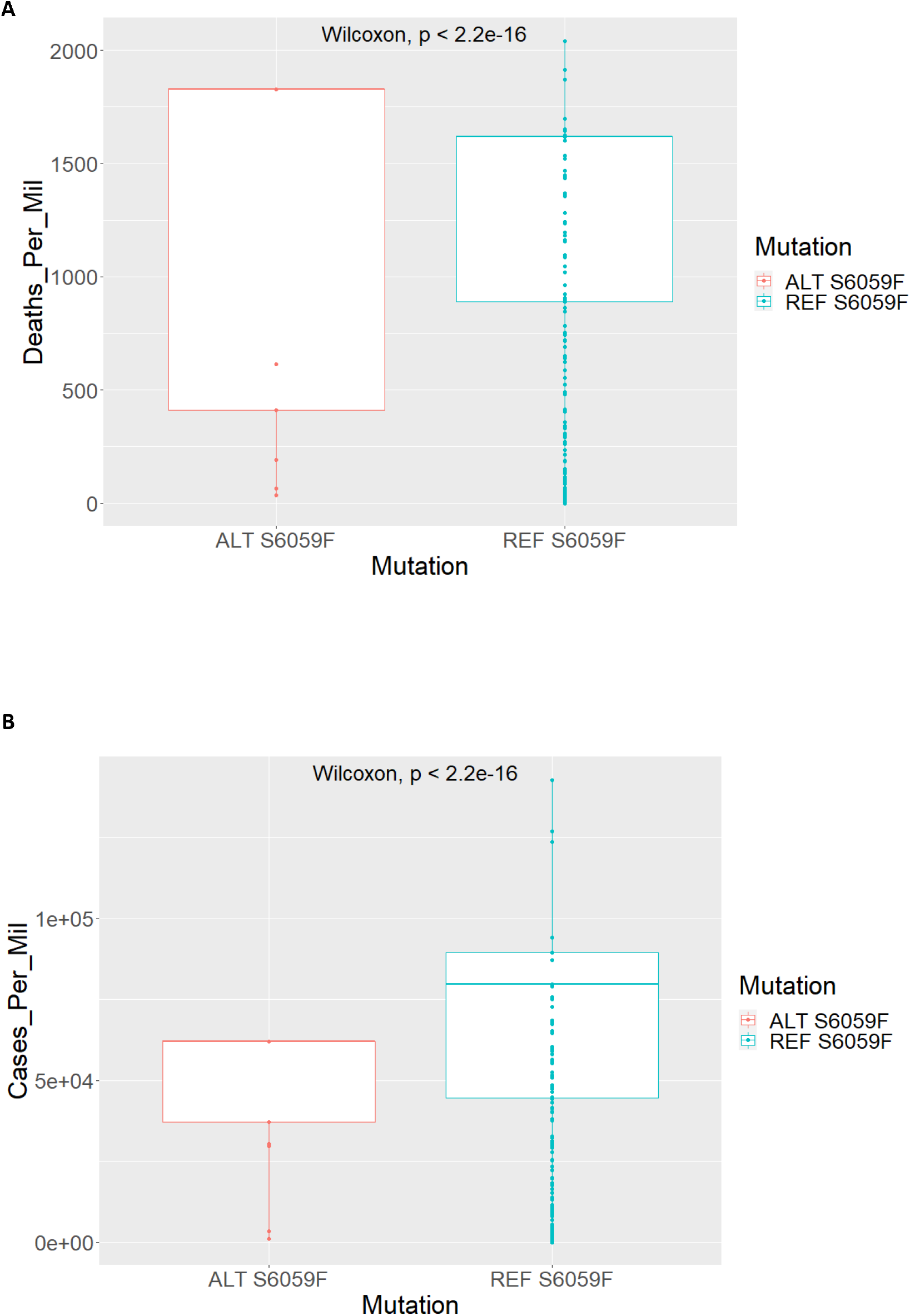
Boxplot distributions for S6059F mutation. **A.** Deaths per million for countries with the ALT and REF forms of the S6059F. **B.** Cases per million for countries with the ALT and REF forms of the S6059F mutation. Note that each data point can represent more than one country.

Applying our previously published glm model [10], on the 18 Cyprus-specific mutations allows for the analysis of these less studied mutations and how they appear to correlate with death and transmission rates. According to the fitting plot of our glm models, the unique S6059F mutation appears to be positively correlated with deaths per million (R=0.61) and neutral with respect to cases (R=0.15) per million (see supplementary **Figure S2**). However, it should be noted that this mutation did not show significance according to the model obtained *p-values* (*p* >0.05). This can most probably be attributed to its low frequency of occurrence across different countries besides Cyprus.

### Structural Analysis

In order to investigate potential reasons why the S6059F mutation appears to be associated with increased mutagenesis, we turn to structural prediction and docking tools. The Ser6059 residue is located on the surface of the NSP14 protein encoded by the *ORF1ab* gene, a bifunctional protein with an N-terminal exonuclease (ExoN) domain with a proofreading function and a C-terminal N7-guanine methyltransferase (N7-MTase) domain, implicated in the methylation of viral RNA cap structures. Both domains are crucial for the maintenance and stability of the viral RNA by lowering its sensitivity to RNA mutagens and evading its degradation from the host immune response [26]. The QHD43415 structure (Estimated TM-score = 0.87) of the NSP14 protein was retrieved from the I-TASSER repository containing the 3D structural models of all proteins encoded by the genome of SARS-CoV-2. The S6059F residue is located in the NSP14 domain directly involved in the physical interaction with NSP10 (residues 1–76 and 119–145), which enhances the ExoN activity by more than 35-fold [26]. The 6059 residue is close to the active site of the enzyme (D90, E92, E191 and D273) [26], and both amino acids are exposed to the surface with RSA values 24.6% and 51.7% for the uncharged Serine (S) and Phenylalanine (F) residues, respectively. Further, in depth molecular analysis reveals disruption of hydrogen bonds and hampering and shifts in other inter-molecular interactions in the ALT-S6059F compared to the REF-S6059F structure of the protein (see **Figure 8A,B**).

**Figure 8.**
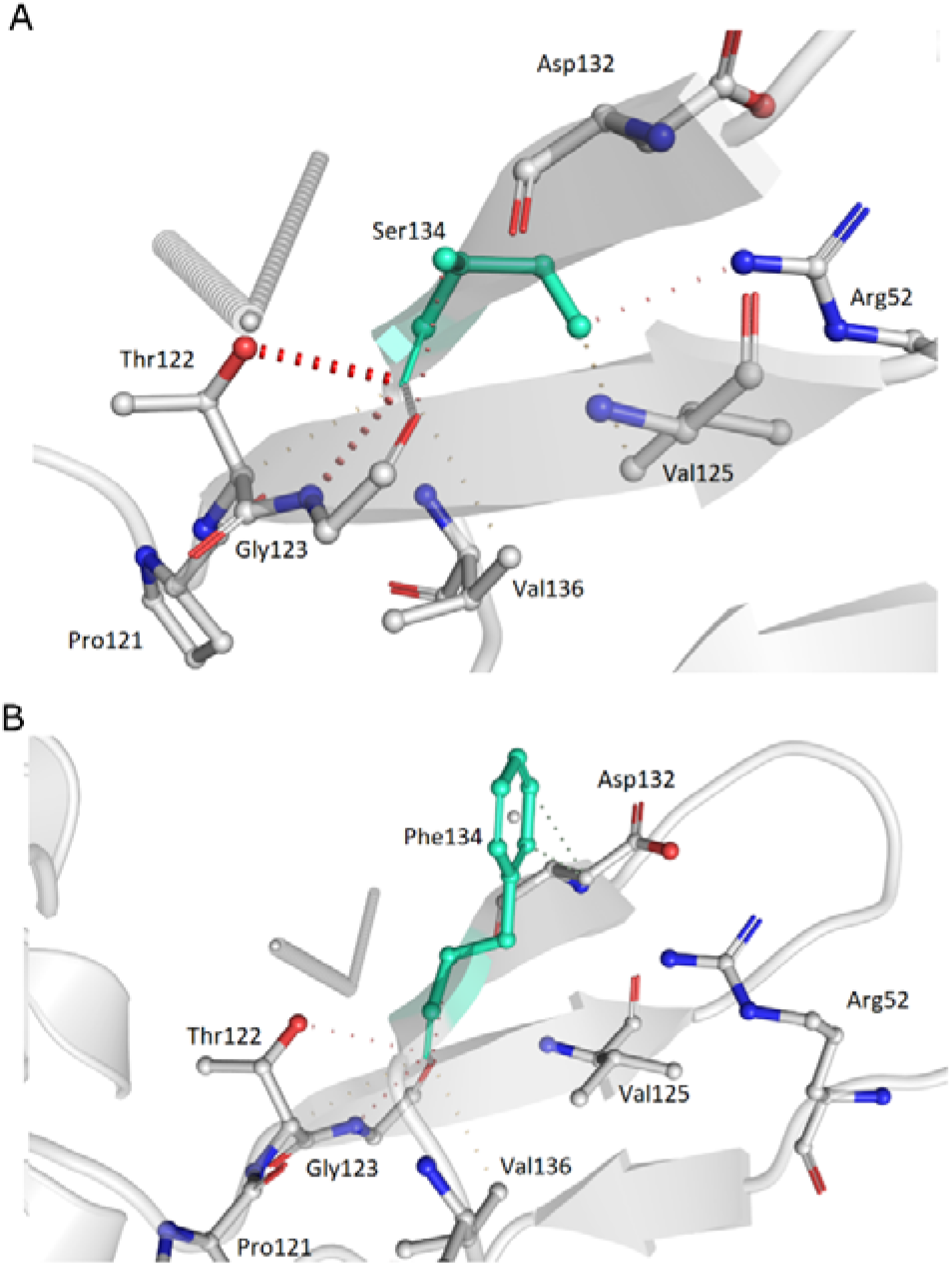
Tertiary and quaternary structural predictions of the NSP14 protein. **A**. Detailed molecular structural conformational changes of NSP14 showing the REF-S6059F. **B**. Same molecular analysis for the structure with ALT-S6059F. Hydrogen bonds in red, weak hydrogen bonds in orange, halogen bonds in blue, ionic interactions in yellow, aromatic contacts in light-blue, hydrophobic contacts in green, carbonyl interactions in pink, VdW interactions in grey.

The experimentally-solved three-dimensional protein structure of the SARS-CoV-2 NSP10 protein was retrieved from the protein databank (PDB ID: 6W75). Protein-protein docking analysis revealed changes on the assembly of NSP14-NSP10 complexes (RMSD 5.841±5.631Å), while the ALT-S6059F-NSP10 complex exhibited a minor increase in complex affinity (ΔG −17.5±0.141kcal/mol and Kd 1.5E-13±2.828E-14M) compared to the REF-S6059F-NSP10 complex (ΔG −16.55±0.495kcal/mol and Kd 8.35E-13±6.576E-13M).

Investigation of the NSP14-RNA interaction was also carried out, using the native 7mer-dsRNA (PDB ID: 4U37) and the derived 7-mer-ssRNA. Upon RNA-protein docking using the HDOCK and MPRDock algorithms, structural changes are evident as demonstrated upon structural alignment for both dsRNA (mean RMSD for both HDOCK and MPRDock 4.255±0.222Å) (**Figure 9A,B**) and ssRNA complexes (mean RMSD for both HDOCK and MPRDock 1.946±2.708Å) (Figure 9C,D).

**Figure 9.**
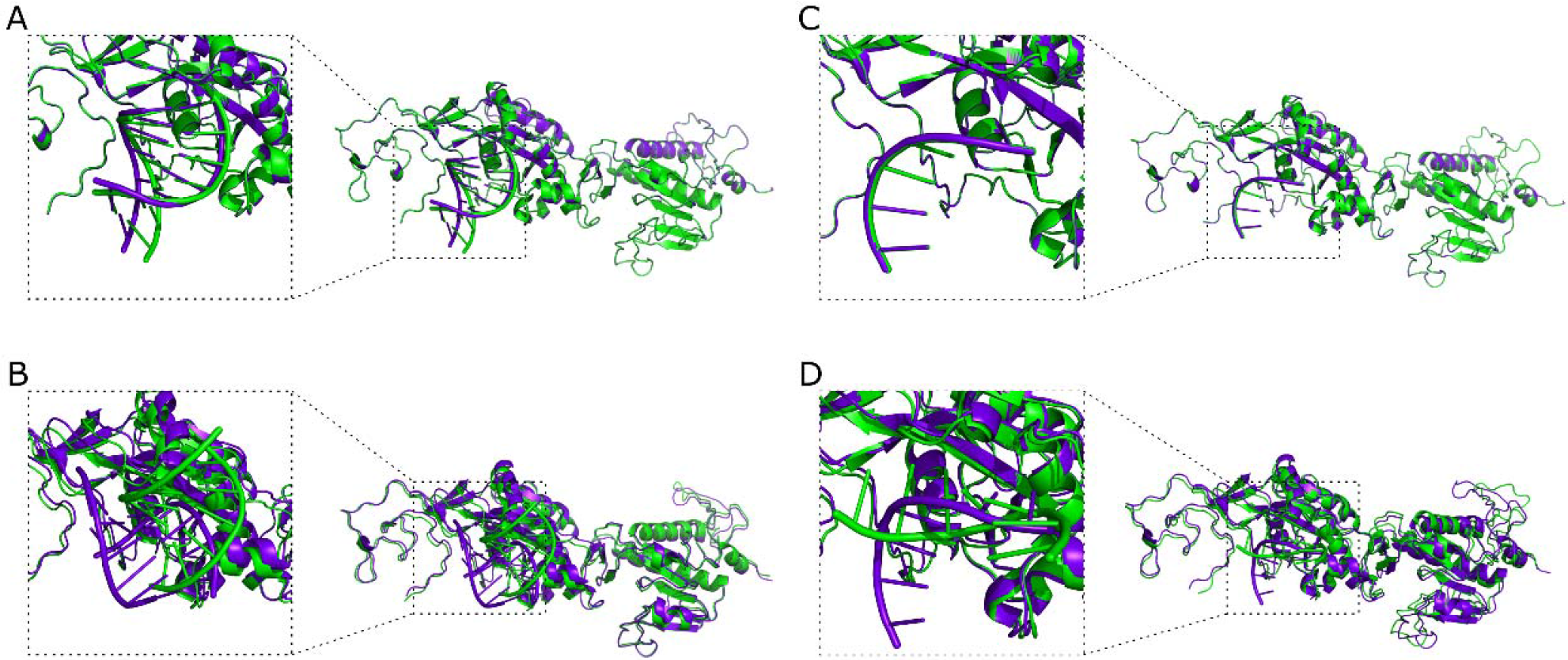
Aligned tertiary and quaternary structural predictions of the complex between the NSP14 protein and RNA (PDB ID: 4U37). **A**. NSP14-dsRNA HDOCK docking simulations’ result. **B**. NSP14-dsRNA MPRDock docking simulations’ results. **C**. NSP14-ssRNA HDOCK docking simulations’ result. **D.** Protein-ssRNA MPRDock docking simulations’ results. REF-S6059F complexes colored in green, ALT-S6059F complexes colored in purple.

Investigation of the NSP10-NSP14-RNA interaction was also carried out. Upon RNA-protein docking using the HDOCK algorithm, structural changes were observed upon structural alignment for both dsRNA (mean RMSD 6.946±4.232Å) (**Figure 10A,B**) and ssRNA complexes (mean RMSD 5.871±5.708Å) (**Figure 10C,D)**.

**Figure 10.**
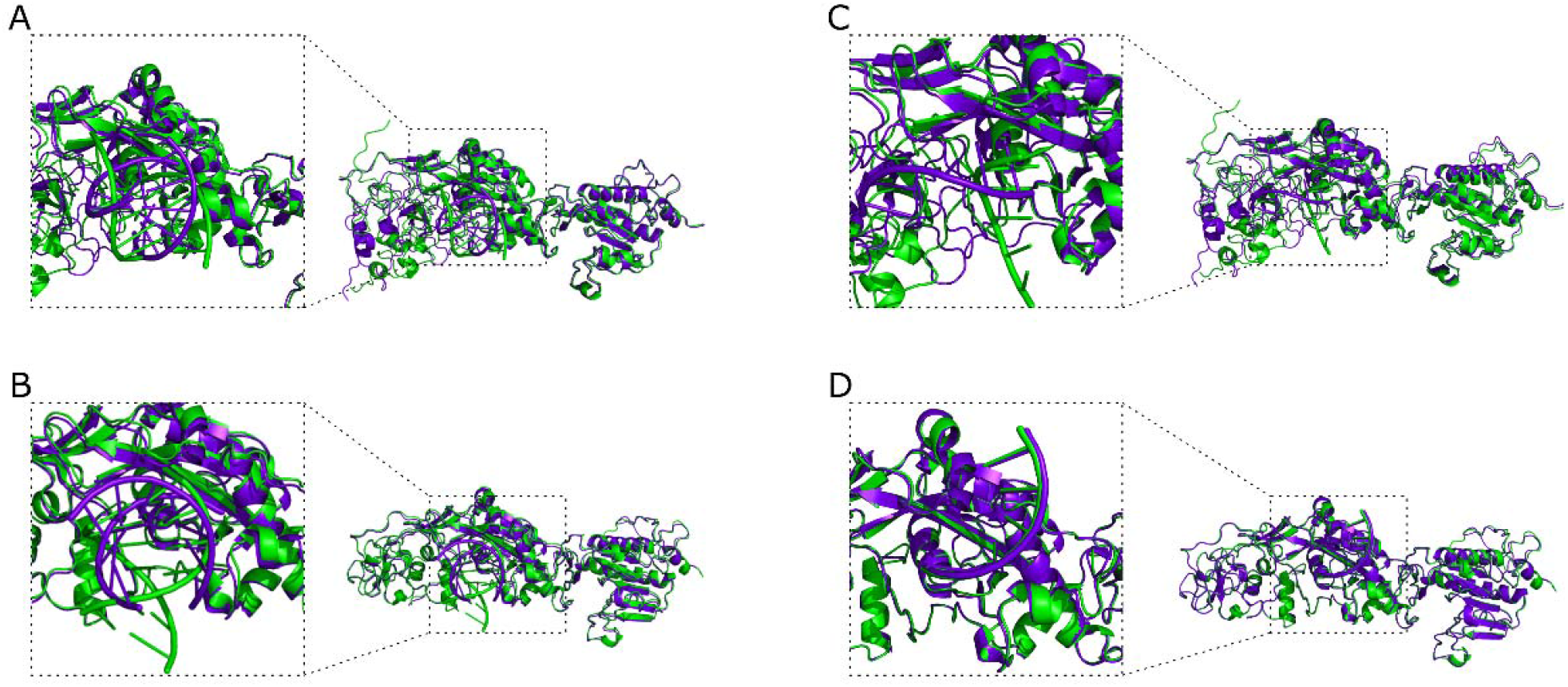
Aligned tertiary and quaternary structural predictions of the complex between the NSP10, NSP14 proteins and RNA (PDB ID: 4U37). **A**. NSP10-NSP14-dsRNA HDOCK docking simulations’ result (using the ClusPro NSP10-NSP14 complex). **B**. NSP10-NSP14-dsRNA HDOCK docking simulations’ result (using the HDOCK NSP10-NSP14 complex). **C**. NSP10-NSP14-ssRNA HDOCK docking simulations’ result (using the ClusPro NSP10-NSP14 complex). **D**. NSP10-NSP14-ssRNA HDOCK docking simulations’ result (using the HDOCK NSP10-NSP14 complex). REF-S6059F complexes colored in green, ALT-S6059F complexes colored in purple.

Since the NSP14 protein is known to be involved in the maintenance and stability of the viral RNA by lowering its sensitivity to RNA mutagens and evading its degradation from the host immune response [26], the large structural changes observed upon mutagenesis at the protein and RNA-protein complex levels, could ultimately lead to a reduced enzyme activity and could be the functional aetiology for observing a greater mutagenesis rate.

Mean free energy values were also reported for the 10 best models attained in order to observe for differences in stability of the NSP14-RNA and the NSP14-NSP10-RNA complexes. HDOCK free energy values are only reported as there were some missing values from the MPRDock algorithm. Results show minor differences in stability between the REF- and ALT-NSP14-RNA complexes for both ds- and ss-RNA molecules examined here (see **Table 2**). More evident differences in complex stability as denoted by free energy scores are observed for the NSP14-NSP10-RNA complexes, with the ALT forms of the complex attaining overall lower free energy values, indicating a higher affinity for RNA binding for the ALT forms of this complex (see Table 2).

**Table 2.**
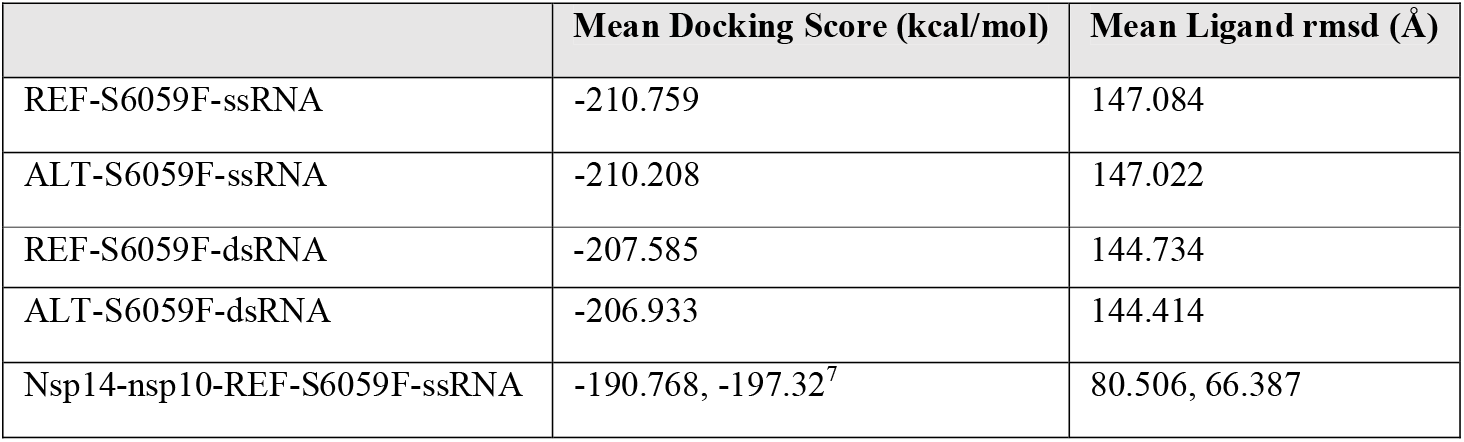
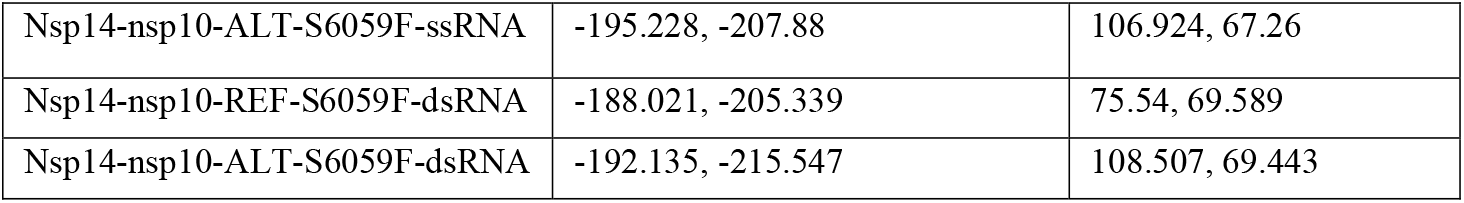
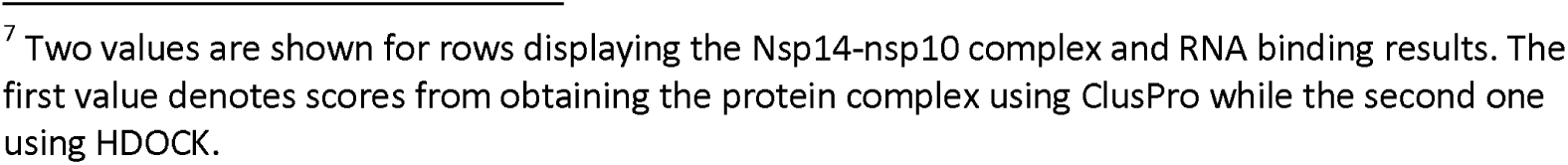
HDOCK Free energy and RMSD scores for REF- and ALT- NSP14-RNA and NSP14-NSP10-RNA complexes.

|  | Mean Docking Score (kcal/mol) | Mean Ligand rmsd (Å) |
| --- | --- | --- |
| REF-S6059F-ssRNA | -210.759 | 147.084 |
| ALT-S6059F-ssRNA | -210.208 | 147.022 |
| REF-S6059F-dsRNA | -207.585 | 144.734 |
| ALT-S6059F-dsRNA | -206.933 | 144.414 |
| Nsp14-nsp10-REF-S6059F-ssRNA | -190.768, -197.32<sup>7</sup> | 80.506, 66.387 |

<sup>7</sup> Two values are shown for rows displaying the Nsp14-nsp10 complex and RNA binding results. The first value denotes scores from obtaining the protein complex using ClusPro while the second one using HDock.

|  |  |  |
| --- | --- | --- |
| Nsp14-nsp10-ALT-S6059F-ssRNA | -195.228, -207.88 | 106.924, 67.26 |
| Nsp14-nsp10-REF-S6059F-dsRNA | -188.021, -205.339 | 75.54, 69.589 |
| Nsp14-nsp10-ALT-S6059F-dsRNA | -192.135, -215.547 | 108.507, 69.443 |

## Discussion

Investigating mutations for SARS-CoV-2 that are unique to a specific country can not only lead to interesting results for the underlying country but also provide insights into viral virulence worldwide. Countries like Cyprus with a small isolated population can be invaluable in assessing viral transmission and death rates, as well as potentially predicting future trends of the virus in the rest of the world. This work was based on the analysis of a small number (N=144) of viral strains from the Cypriot population. To this end we initially perform basic phylogenetic and lineage analysis as previously reported for these samples [9]. We extend previous work by selecting for 18 Cyprus-specific mutations exhibiting a high frequency of occurrence that is unique to Cyprus compared to the rest of the world. We highlight a single mutation with the highest frequency of occurrence in Cyprus alone and further analyse this by mutation tracking and regressions analysis (glms) in order to obtain a greater understanding of the nature of this mutation. We show that this mutation causes an amino acid change (S6059F) on the NSP14 exoribonuclease of the *ORF1ab* protein. We furthermore assess whether this mutation affects the molecular functionality of the NSP14 protein, which is known to be implicated in viral mutation proofreading during replication. We provide evidence that it may actually allude to a dysfunctional NSP14 protein that causes hypermutability in the strains with this mutation. This is further supported by structural modelling of the NSP14 protein with and without the S6059F mutation, which clearly points to a different structural conformation of the ALT vs. the REF form of the protein. This structural variation is exhibited by the NSP14 protein alone as well as in complex with NSP10 and with both ds- and ss-RNA molecules. Moreover, free energy values report greater stability in the ALT-NSP14-NSP10-RNA complex. The mutation-generated alteration in RNA or DNA affinity has also been investigated with other viral proteins that are involved in RNA/DNA processing during viral replication. A recent example comes from the Herpes Simplex Virus UL42 protein which binds DNA and plays an essential role in viral DNA replication by acting as the polymerase accessory subunit. It has been shown that engineered viruses expressing mutant forms of the UL42 proteins that increase its affinity for DNA binding, exhibited increased mutation frequencies and elevated ratios of virion DNA copies [34]. These results suggest the SARS-CoV-2 S6059F-generated increased affinity for RNA seen in our structural models, may be the cause of the hypermutation also observed in our data for the Cypriot strains with the ALT form of the S6059F mutation. As previously reported, certain viruses may have evolved so that their RNA or DNA binding proteins, implicated in viral replication, neither bind DNA too tightly nor too weakly to optimize virus production and replication [34]. It is important to stress that our proposed mechanism for the S6059F mutation alluding to a dysfunctional NSP14 protein, which in turn may cause hypermutability, requires additional *in-vitro* experimental verification before it can be fully validated.

We propose a theory whereby the increased mutability caused by decreased proofreading as a consequence of the S6059F mutation, causes a greater diversity of viral strains or quasispecies resident within the same host. This can potentially impact the infection patterns descending from this host by altering transmission rates and possibly death rates. We outline two potential scenarios where this could be of significant impact for downstream viral infection trends. If the “main” nested strain is of high virulence, viral replication with increased mutability will allude to a greater range of viral quasispecies from which to infect downstream hosts, thus less chances of the host transmitting the strain with high virulence. If, on the other hand, the nested strain is one of low virulence, increased mutability could potentially lead to the development and consequent transmission of more virulent strains in downstream infections. One very important aspect that ought to be discussed and investigated further, is the potential impact of such viable mutations that cause hypermutability on vaccination efficacy. It can be argued that infections caused from viral strains with such proofreading dysfunction can act as generators of diverse viral strains that allude current vaccination attempts at a faster rate compared to strains with more regular proofreading functionality.

