## Supplementary Information for "In depth analysis of Cyprus-specific mutations of SARS-CoV-2 strains using computational approaches"

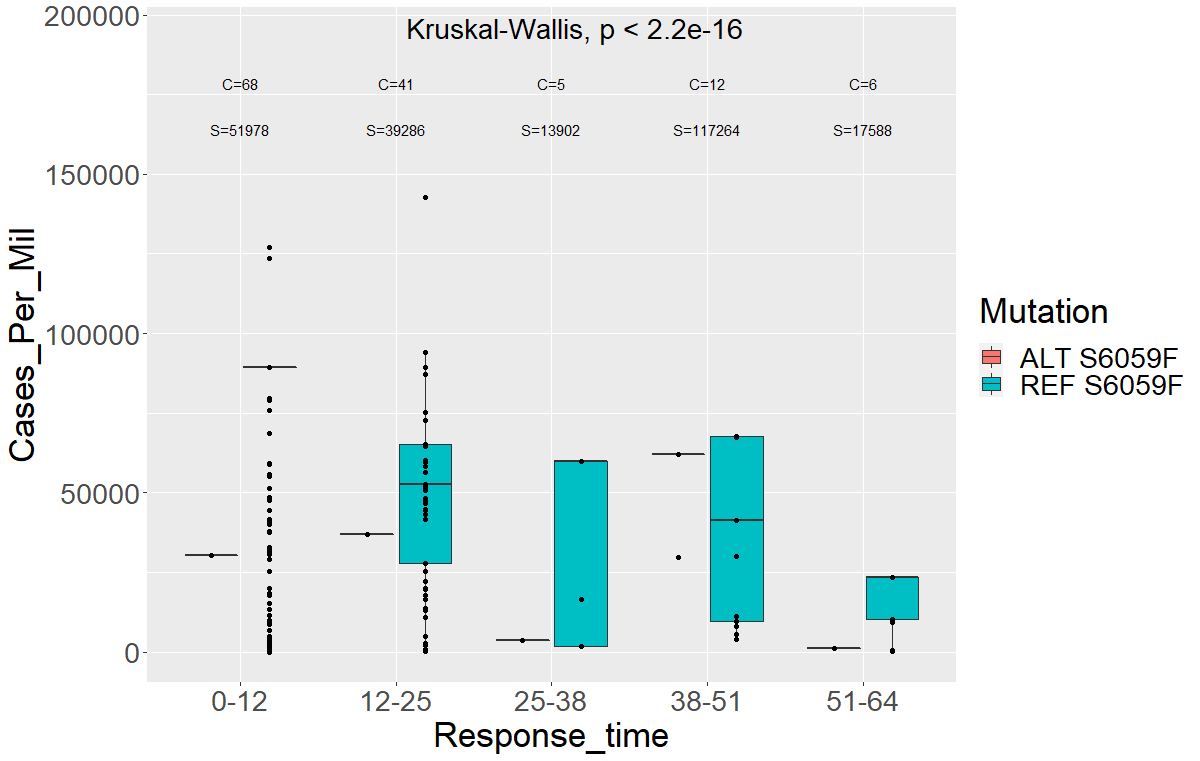

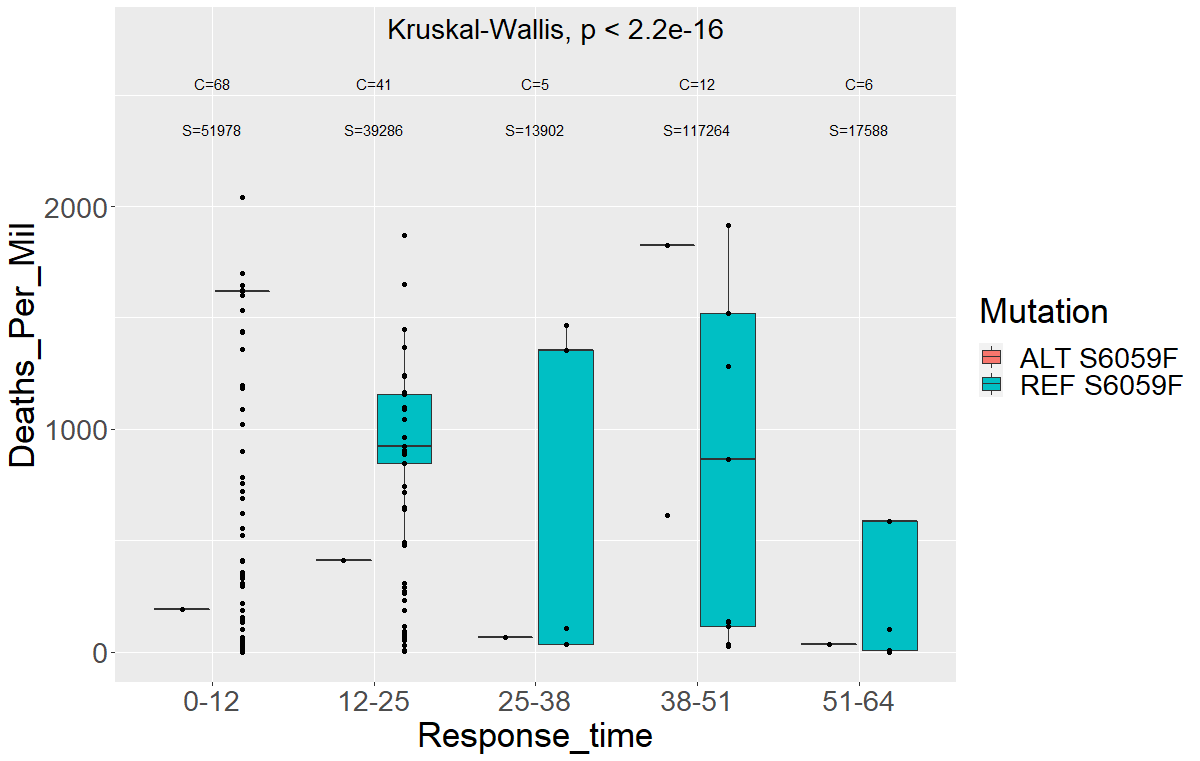

**A**

**B**

**Figure S1 A.** Deaths per million for countries with the S6059F mutation and the reference mutation including response time separation. *C* denotes the number of unique countries in the group and *S* is the number of strains in the group. **B.** Cases per million for countries with the S6059F mutation and the reference mutation including response time separation. *C* and *S* are as denoted for panel **B**.

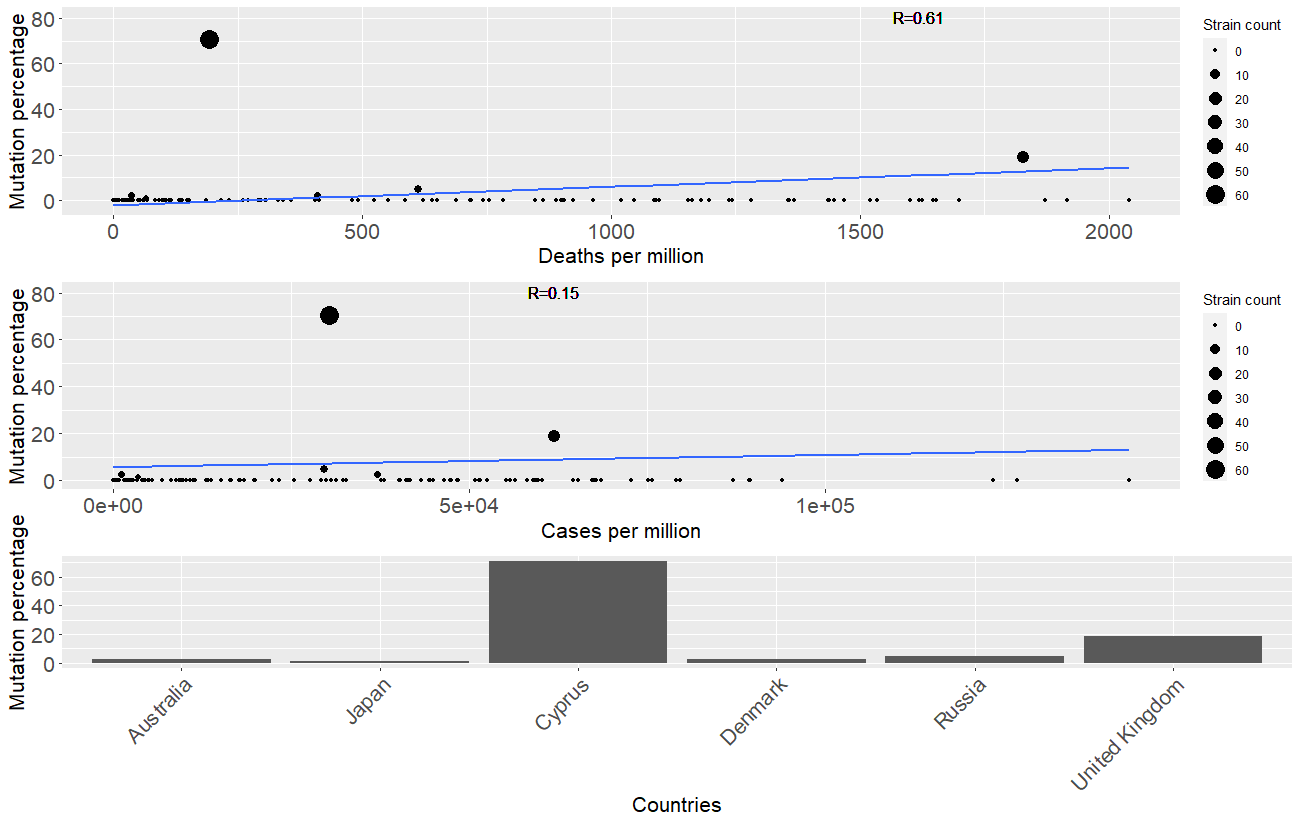

**A**

**B**

**C**

**Figure S2.** **Analyses plots for the ORF1ab protein mutation at position S6059F** **A**. Regression model line showing the simplified fit for mutations’ percentage across countries and the death rate per million for each country. Pearson’s correlation is shown by the *R* value. **B**. Similar regression fit for mutations percentage across countries this time showing cases per million for each country. **C**. Detailed histogram of the percentage occurrence of the mutation across different countries. Countries are sorted with increasing deaths per million.
